# Taurine pangenome uncovers a segmental duplication upstream of *KIT* associated with depigmentation in white-headed cattle

**DOI:** 10.1101/2024.02.02.578587

**Authors:** Sotiria Milia, Alexander S. Leonard, Xena Marie Mapel, Sandra Milena Bernal Ulloa, Cord Drögemüller, Hubert Pausch

## Abstract

Cattle have been selectively bred for coat color, spotting, and depigmentation patterns. The assumed autosomal dominant inherited genetic variants underlying the characteristic white head of Fleckvieh, Simmental, and Hereford cattle have not been identified yet, although the contribution of structural variation upstream the *KIT* gene has been proposed. Here, we construct a graph pangenome from 24 haplotype assemblies representing seven taurine cattle breeds to identify and characterize the white head-associated locus for the first time based on long-read sequencing data and pangenome analyses. We introduce a pangenome-wide association mapping approach which examines assembly path similarities within the graph to reveal an association between two most likely serial alleles of a complex structural variant 66 kb upstream *KIT* and facial depigmentation. The complex structural variant contains a variable number of tandemly duplicated 14.3 kb repeats, consisting of LTRs, LINEs, and other repetitive elements, leading to misleading alignments of short and long reads when using a linear reference. We align 250 short-read sequencing samples spanning 15 cattle breeds to the pangenome graph, further validating that the alleles of the structural variant segregate with head depigmentation. We estimate an increased count of repeats in Hereford relative to Simmental and other white-headed cattle breeds from the graph alignment coverage, suggesting a large under-assembly in the current Hereford-based cattle reference genome which had fewer copies. We show that exploiting assembly path similarities within graph pangenomes can reveal trait-associated complex structural variants.

## Introduction

Natural selection, domestication, and selective breeding have shaped the genetic and phenotypic variation of modern cattle (*Bos taurus*). Hundreds of cattle breeds are recognized worldwide belonging to the indicine (*Bos taurus indicus*) or taurine (*Bos taurus taurus*) subspecies (Loftus et al., 1994). Coat color, spotting, and piebald patterns differentiate the breeds as these traits have frequently been used for breed formation (Cieslak et al., 2011). Depigmentation phenotypes can vary from tiny white spots, white head and leg markings, large irregular white patches, symmetrical patterns such as a white band around the midsection or white stripes along the dorsal and ventral midlines, to almost complete depigmentation in white born animals (Olson, 1981). The genetic basis of coat color variation has been studied extensively in many domesticated species including cattle, and several large-effect loci, sometimes associated with pleiotropic effects, have been identified (Bannasch et al., 2021; Durkin et al., 2012; Haase et al., 2009; Henkel et al., 2019; Joerg et al., 1996; Rubin et al., 2012; Trigo et al., 2021). Variation in coat color is mostly the result of spontaneous mutations, and the derived trait-associated alleles are often propagated through selective breeding because the mutant animals’ striking appearance and special value to their owners result in iconic, breed-defining phenotypes. In particular, allelic variation nearby *KIT* encoding KIT proto-oncogene, receptor tyrosine kinase is responsible for several breed-defining coat color patterns (Durkin et al., 2012; Grosz and MacNeil, 1999; Hayes et al., 2010; Küttel et al., 2019; Liu et al., 2009; Qanbari et al., 2014).

Structural variants (SVs) affecting the expression of *KIT* contribute to depigmentation phenotypes in cattle and other species (Artesi et al., 2020; Dürig et al., 2017; Durkin et al., 2012; Giuffra et al., 2002; Küttel et al., 2019; Nagle et al., 1995; Venhoranta et al., 2013). A series of three *KIT*-related alleles, all complex structural rearrangements expected to reflect gain of function variants resulting in dysregulated *KIT* expression, cause the color-sided pattern in cattle and yak (Artesi et al., 2020; Durkin et al., 2012; Küttel et al., 2019). A sequence-based genome-wide association study mapped the white-headed phenotype in Fleckvieh cattle also to a region encompassing *KIT* (Qanbari et al., 2014), but the causal variant and functional mechanism remain unclear. Increased sequencing coverage upstream of *KIT* has been detected in short-read sequenced Hereford and Simmental cattle, both of which have a characteristic white head (Whitacre, 2014), suggesting that a copy number variant might contribute to this enigmatic phenotype. Due to the difficulty of resolving SVs using short reads and alignment-based methods (Bickhart and Liu, 2014), further characterization and validation of this region has not been attempted to date.

Long-read sequencing and genome assembly methods enable detecting and resolving SVs that were previously difficult to identify using short-read sequencing data. Pangenomes, which incorporate a series of assemblies rather than a single reference genome, can fairly represent all types of variation present in a population. There are several approaches to constructing pangenomes, using reference-backed (Li et al., 2020), phylogeny-guided (Armstrong et al., 2020), or all-to-all (Garrison et al., 2023) alignments. These graph pangenomes reduce reference bias because the read aligners can “see” all alleles simultaneously, improving variant calling accuracy and enabling SV genotyping (Liao et al., 2023). Some graph aligners, like vg giraffe (Sirén et al., 2021) and experimental updates to GraphAligner (Rautiainen and Marschall, 2020), can exploit haplotype information stored in the graph (which input assemblies took which variant paths), further improving genotyping speed and accuracy.

Here, we combine 14 newly assembled HiFi-based haplotypes with ten previously published assemblies to create a 24-assembly taurine pangenome graph. We introduce a novel mapping approach to identify candidate regions for the white-headed phenotype directly from the graph structure, and then align 250 short-read samples from 15 breeds to confirm the association between serial alleles of a complex structural variant upstream of *KIT* and the phenotype.

## Results

### Head pigmentation segregates among sequenced breeds

We analyzed 24 long read sequencing-based assemblies representing seven taurine cattle breeds to investigate the genetic underpinnings of head pigmentation. These assemblies included 14 newly generated HiFi-based haplotypes from Simmental, Evolèner, Rätisches Grauvieh, Brown Swiss, Original Braunvieh, and Braunvieh cattle that were assembled through trio-binning (N=6) or a dual-assembly approach (N=8) when parental data were not available (Table 1). The newly generated assemblies were highly contiguous with a mean contig N50 of 78.2 Mb, as well as highly accurate with a mean quality value (QV) of 48, or approximately 1 error every 62 kb. We also validated the gene completeness with compleasm, identifying on average 98% of expected conserved genes. We observed variation in the assembly quality even for samples with comparable sequencing coverage. However, this could mostly be explained by the sex of the haplotype (the cattle X chromosome is roughly three times longer than the cattle Y and has ∼350 additional conserved genes), variation in read length/quality (Liao et al., 2023), and trio versus dual haplotype phasing.

**Table 1.** Fourteen newly HiFi-assembled taurine haplotypes from Evolèner (EVO), Simmental (SIM), Rätisches Grauvieh (RGV), Original Braunvieh (OBV), or Brown Swiss (BSW) ancestry that were created with trio-binning (Trio) or a dual-assembly (Dual) approach. Braunvieh (BV) indicates samples with both Brown Swiss and Original Braunvieh ancestry. HiFi coverage is given in gigabases per animal, and so is shared when both haplotypes are given. N50 is calculated with respect to the ARS-UCD1.2 size. The quality value (QV) score is estimated with merqury from short reads. Gene completeness is calculated by compleasm on the cetartiodactyl ODB10 gene set. * indicates male triobinned haplotypes including only a Y chromosome (no X chromosome), which can result in small genome size, lower N50, reduced gene completeness etc. ^†^ indicates male F1s that were assembled without parental information, and so X and Y chromosomes may be split between the dual haplotypes.

| Sample / Acronym | HiFi Coverage (Gb) | Haplotype | Size | contigs | N50 | QV | Gene completeness |
| --- | --- | --- | --- | --- | --- | --- | --- |
| GIRxSIM / SIM_2 | 91.4 | Trio | 3.16 | 886 | 52.5 | 53.1 | 99.6 |
| RGVxSIM / SIM_3 | 140.1 | Trio | 3.21 | 682 | 89.9 | 58.8 | 99.5 |
| RGVxSIM / RGV* |  |  | 3.09 | 816 | 91.9 | 58.3 | 97.2 |
| DWZxEVO / EVO | 139.1 | Trio | 3.19 | 527 | 92.1 | 57.5 | 99.7 |
| Brown Swiss / BSW_3* | 111.8 | Trio | 3.11 | 1031 | 80.6 | 46.7 | 98.4 |
| Brown Swiss / BSW_4 |  |  | 3.11 | 857 | 71.2 | 47.1 | 98.3 |
| Original Braunvieh / OBV_3 | 135.8 | Dual† | 2.95 | 2250 | 72.6 | 44.5 | 97.0 |
| Original Braunvieh / OBV_4 |  |  | 2.99 | 1550 | 71.5 | 44.9 | 97.9 |
| Braunvieh / BV_1 | 122.9 | Dual† | 2.95 | 2336 | 74.9 | 47.4 | 98.2 |
| Braunvieh / BV_2 |  |  | 2.99 | 1755 | 72.4 | 47.6 | 96.6 |
| Braunvieh / BV_3 | 145.0 | Dual† | 3.02 | 1701 | 83.2 | 43.1 | 95.1 |
| Braunvieh / BV_4 |  |  | 2.85 | 2099 | 82.6 | 44.2 | 96.6 |
| Braunvieh / BV_5 | 131.0 | Dual† | 3.05 | 1373 | 85.0 | 38.2 | 98.6 |
| Braunvieh / BV_6 |  |  | 2.91 | 1681 | 74.6 | 39.1 | 96.5 |

A Simmental haplotype (SIM_2) produced through trio-binning required some manual curation to correct an under assembled region of interest introduced later (see Methods). We further collected nine publicly available long read-based haplotype assemblies including a Simmental (Heaton et al., 2021) and a Highland (Rice et al., 2020) haplotype assembled with ONT reads, as well as seven HiFi-assembled Brown Swiss and Original Braunvieh haplotypes (Leonard et al., 2023a, 2023b, 2022) (Supplementary Table 1). We also included the cattle reference genome (ARS-UCD1.2) which was assembled from PacBio Continuous Long Read data collected from a Hereford cow (Rosen et al., 2020).

Cave paintings indicate that European aurochs, the wild ancestor of taurine cattle, had a colored head (Stokstad, 2015) suggesting that the white head seen in several breeds emerged as the result of mutations that occurred during or after domestication. A white head (Figure 1A) is characteristic for Simmental and Hereford cattle while cattle from the other five breeds included in the pangenome have almost fully pigmented heads. The head pigmentation of all individuals (or their parents for the trio-binned assemblies) from which the haplotypes were assembled matched these expectations. An approximate tree constructed with mash from the assemblies (Figure 1B) reveals that Simmental and Hereford belong to different clades despite sharing the white-headed phenotype, while the overall tree is compatible with previously reported breed phylogenies (Decker et al., 2014). Simmental cattle have Central European ancestry while Hereford originate from the British Isles (Decker et al., 2014, 2009). However, it remains unclear whether the white head-associated allele arose independently in these two breeds or if it was introgressed from one to the other.

**Figure 1.**
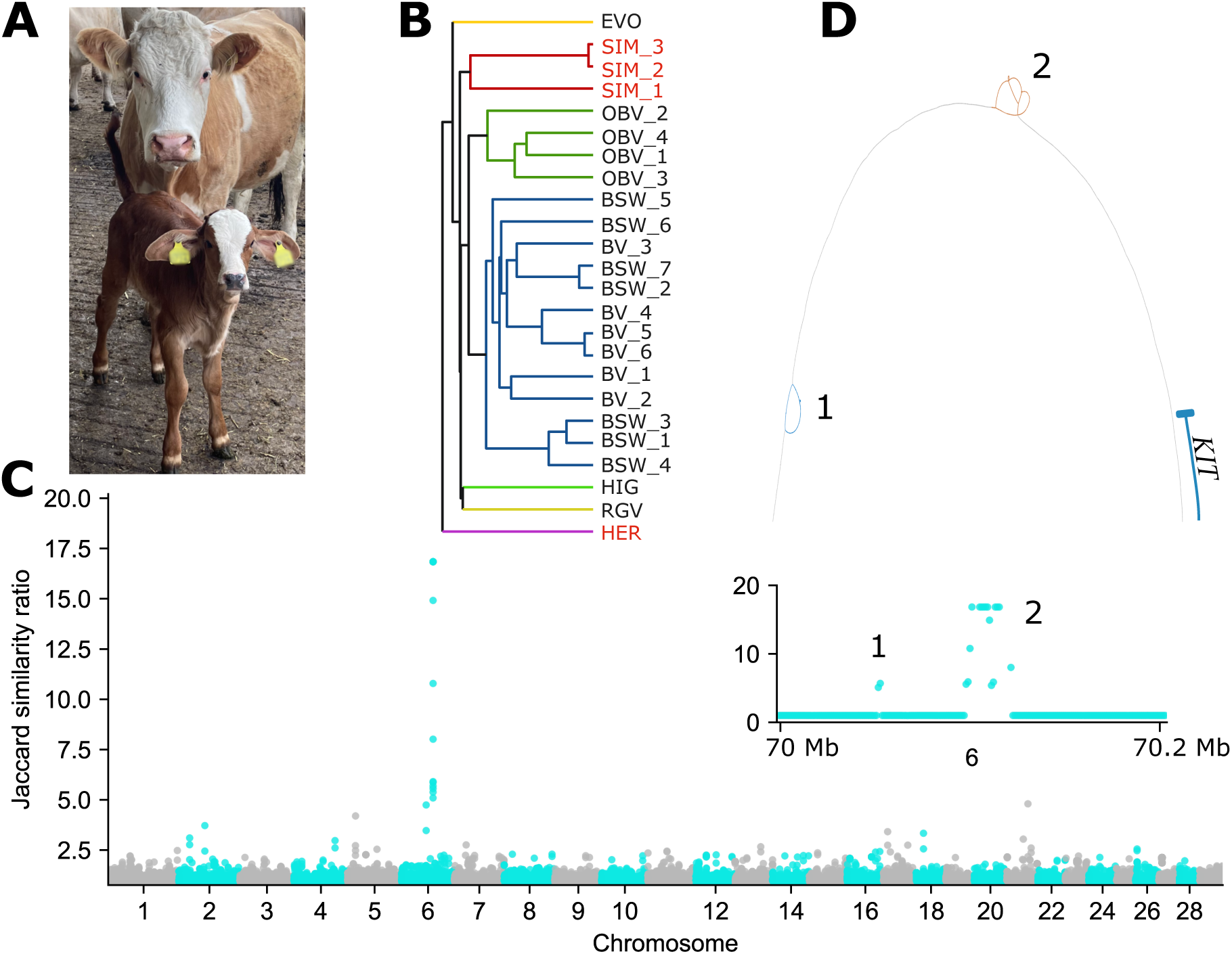
White head in taurine cattle is associated with an SV bubble on chromosome 6. (A) Purebred white-headed Simmental cow with its daughter showing a large white blaze. The daughter originates from a cross with a Gir sire. (B) Mash-based phylogenetic tree of the 24 input taurine assemblies across chromosome 1. Sample names from breeds with a white/color-headed phenotype are red/black respectively, while each breed cluster on the tree has its own color. (C) Jaccard similarity ratio within 1 kb bins (with respect to the ARS-UCD1.2 reference genome) of the pangenome graph between white-headed and color-headed groups. There were two separate regions of interest, both on chromosome 6 upstream of KIT (indicated with 1 and 2 in the inset). (D) An approximately 150 kb region containing the two regions of interest and the first coding exon of the KIT gene.

The 1000 Bull Genomes Project generated a substantial amount of short-read sequencing data for three breeds (Simmental, Hereford, and Braunvieh) included in our pangenome. Two of these breeds exhibit white heads, while one has a colored head. Signature of selection analyses using these short-read sequencing-derived variants revealed a strong selective sweep near *KIT* in both white-headed breeds (Supplementary Figure 1). This finding corroborates previous mapping attempts and suggests a shared genetic origin of the white-headed phenotype (Fontanesi et al., 2010; Qanbari et al., 2014; Whitacre, 2014). The strongest sweep signal was upstream of *KIT* supporting that the underlying mutation has a regulatory role. Regulatory SVs impact the expression of *KIT* in various species (Berrozpe et al., 2006; Brooks et al., 2008; Durkin et al., 2012; Giuffra et al., 2002; Henkel et al., 2019), leading us to hypothesize that an SV affecting regulatory sequence also underlies head depigmentation in Simmental and Hereford cattle.

### Pangenome graph construction and association testing identifies a trait-associated complex structural variant

Genome-wide association studies for head pigmentation in cattle have not yet considered SVs due to the limitations of short-read sequencing technology used in previous genotyping efforts. To make complex SVs amenable to association testing, we incorporated the 24 haplotype assemblies into a graph pangenome. We constructed graph pangenomes for each autosome separately for the 24 taurine assemblies using reference-free base-level alignment with PGGB (Garrison et al., 2023). Graph construction required approximately 484 CPU hours in total across the autosomes with a peak memory usage of 24.5 Gb. Combined, the graphs contained 3,701,244,547 nucleotide bases across 57,964,174 nodes connected by 79,388,152 edges. The reference genome autosomes only cover 2,489,368,269 bases, leading to approximately 1,212 Mb of non-reference sequence. Much of this non-reference sequence originates from centromeres assembled on the HiFi assemblies. The graphs contained 1,145 nodes larger than 100 kb, totaling 816 Mb of sequence. Of these, 695 nodes contained 588 Mb of sequence annotated as centromeric satellites, as expected given the challenges in aligning and collapsing homologous centromeric sequence in graph building (Liao et al., 2023). There were also several large, non-repetitive nodes that were erroneously not merged into syntenic regions in the graphs, inflating the amount of non-reference sequence. The largest instance was a single 2.6 Mb node on BTA10:22720789-25310574, where the actual coordinates with poor alignments are between approximately 23-24 Mb (Supplementary Figure 2). However, this specific region in ARS-UCD1.2 has previously been reported for poor alignment and imputation quality (Pausch et al., 2017), and so this issue is not unique to graph building. These artefactually large nodes are overall rare, and do not substantially impact graph-wide analyses.

We searched for regions in the graph pangenome where the haplotype paths were classified into distinct groups (white-headed Simmental and Hereford vs. all others with colored heads) to identify variants associated with a white head. To do so, we assessed the pairwise Jaccard similarity over the paths taken by each assembly through 1 kb windows and took the ratio of average similarities between similar phenotypes and opposite phenotypes. A large ratio indicates regions of the graph where e.g., a Simmental and Hereford assembly are similar, a Brown Swiss and Highland are similar, but the Simmental and Brown Swiss are distinct. This “phenotype signature”-approach identified a single prominent peak at BTA6:70.09-70.12 Mb, with a second, less strongly associated cluster very close by at BTA6:70.05 Mb (Figure 1C). Several smaller, single window peaks (e.g., BTA6:54.7 Mb and BTA21:47.3 Mb) were primarily driven by most likely breed-specific SVs of Simmental cattle (Supplementary Figure 3), and so were not considered for subsequent analysis.

The most prominent association signal was from a large, complex bubble spanning BTA6:70099582-70123136. All 20 assemblies from color-headed animals effectively contained a 20.6 kb deletion with respect to the reference genome, while the three white-headed Simmental assemblies contained more sequence than the Hereford-based ARS-UCD1.2 assembly. A less prominently associated second cluster was a 6.8 kb insertion present approximately 47 kb further upstream in color-headed animals of both breeds (SIM and HER), in addition to these assemblies traversing the ∼1 kb sequence contained in BTA6:70052702-70053557 twice. The white-headed animals only traversed those reference coordinates once. These two bubbles were respectively 66 and 113 kb upstream the translation start site of *KIT* (NM_001166484.1) at 70,166,794 bp (Figure 1D), coinciding with the strong signatures of selection detected from short read-derived variant genotypes of Simmental and Hereford cattle (Supplementary Figure 1B).

### A graph bubble containing repetitive sequence is associated with coat color of the bovine head

We characterized the sequence content within and adjacent to the most strongly trait-associated graph bubble to reveal a possible functional explanation for the depigmented heads of Simmental and Hereford cattle. The bubble contains a total number of 276 nodes which correspond to a combined length of ∼43 kb for each of the three Simmental haplotypes and ∼26 kb for the Hereford-based ARS-UCD1.2 assembly. The three Simmental assemblies contain three copies of a 14.3 kb sequence, which form the bubble. The ARS-UCD1.2 reference genome contains two copies of this segment but misses 1.8 kb and 800 bp of sequence from each copy respectively relative to the full 14.3 kb sequence observed in the Simmental assemblies. The assemblies of the color-headed cattle appear to have only part (∼5.5 kb) of one copy of the 14.3 kb sequence, corresponding to a deletion of 20.6 kb relative to the ARS-UCD1.2 reference genome. The summed length of the nodes within the graph bubble is approximately 29 kb, as some nodes are traversed multiple times across the tandemly duplicated sequences, demonstrating there is sufficient sequence similarity between and across haplotypes to share and reuse nodes (Figure 2A).

**Figure 2.**
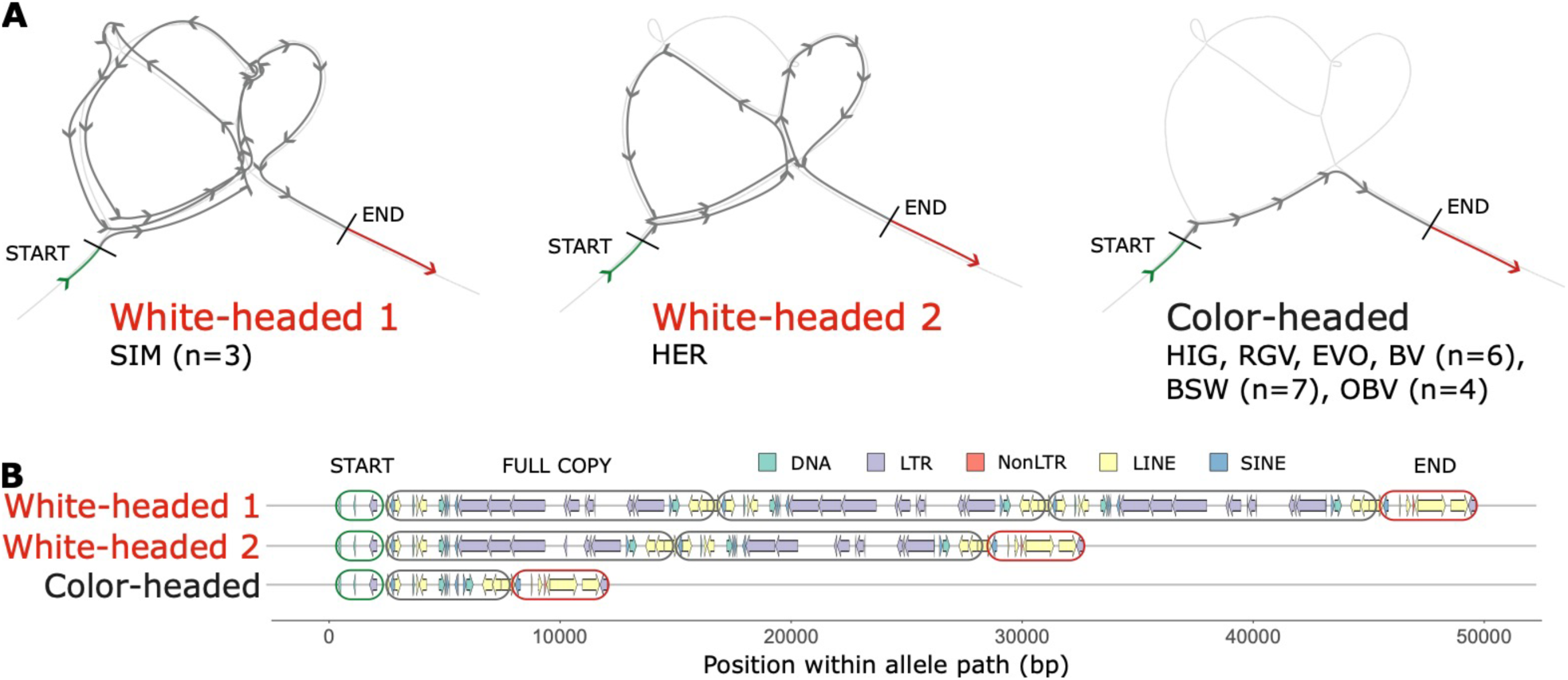
Topology of a trait-associated bubble upstream the KIT gene. (A) Bandage plots showing the paths traversed for the three observed alleles (two for white-headed breeds and one for color-headed) through the pangenome. Unless stated otherwise, n=1 for the breed. The grey arrow paths indicate the segmental duplication regions, while the green and red arrow paths respectively indicate the sequence preceding and following the segmental duplication. (B) Repeat structure within the candidate SV region, for 5 classes of genomic elements. The start and end sequences are taken from (A), while the full copy refers to the 14.3 kb segmental duplication copy found in all three Simmental assemblies. The arrow direction denotes the strand of the repeat element (+/forward and -/reverse).

The full sequence of the segmental duplication identified in the Simmental assemblies contains transposable elements consisting mostly of long terminal repeats (LTRs). The structure of each copy of the segmental duplication starts with a short interspersed nuclear element (SINE) and ends with a long interspersed nuclear element (LINE) (Figure 2B). This repetitive nature of the duplication contributes to the complexity and variability of the region, thus leading to misleading alignments of and subsequently variant calls from both short and long reads when using a linear reference (Supplementary Figure 4, Supplementary Figure 5). The color-headed animals share the sequence immediately preceding and following the segmental duplication but have a truncated copy of the tandemly duplicated sequence that misses most of the LTR elements.

A detailed functional investigation and identification of putative *cis*-regulatory sequences within or nearby the segmental duplication is challenging due to the lack of a comprehensive functional annotation and three-dimensional structure of the bovine genome. Therefore, we characterized the sequence content of the trait-associated region based on human regulatory elements for which we lifted the coordinates onto the ARS-UCD1.2 cattle reference genome. We mapped human promoters, enhancers, and transcription factor binding sites to the orthologous region of bovine ARS-UCD1.2 identifying various regulatory elements nearby the segmental duplication (Supplementary Figure 5). We mapped four cis-regulatory elements from human ENCODE including three distal enhancer-like sequences within a 50 kb window of the insertion using LiftOver. Two of these (EH38D3577379 predicted at 6:70063721-70064026 and EH38D3577384 predicted at 6:70065620-70065838) also overlapped permissive enhancer sequences from the FANTOM5 regulatory annotations of the human genome. These sequences are approximately 103 kb upstream of the translation start site of *KIT*. There were no regulatory elements predicted within the sequence that is tandemly duplicated in white-headed animals, nor were any predicted regulatory elements disrupted by the duplication.

### Pangenome-based genotyping in a diversity panel of short-read sequenced cattle supports that a complex SVs underlies depigmentation in white-headed cattle

To further validate an association between the segmental duplication and the white-headed phenotype, we collected 250 publicly available whole genome short read sequencing samples spanning 15 taurine breeds (Supplementary Table 2). The average read depth of these samples estimated from alignments against ARS-UCD1.2 was 12.3 (± 4.4)-fold. Cattle of the Simmental, Hereford, Groningen White Headed, Kazakh White Headed, Yaroslavl, Montbéliarde, and Normande breeds show different types of the white-head phenotype, while the other breeds do not (although some are completely white colored or have minor white head markings). We validated the breed-identification associated with each sample through a principal component analysis of sequence variant genotypes called from reference alignments (Supplementary Figure 6), with breeds clustering largely as expected. Cattle from white-headed and color-headed breeds were sometimes more closely clustered than two cattle from white-headed breeds (as observed earlier in Figure 1B), indicating that the allele is likely not private to a particular ancestral taurine lineage but is segregating in more distantly related breeds, possibly indicating admixture. Otherwise, this could also indicate that the causal mutation event occurred before the formation of modern cattle breeds, or allelic heterogeneity.

We then aligned all short-read sequencing samples with vg giraffe to a pangenome subgraph spanning BTA6:69099582-71123136 (1 Mb up- and downstream of the most significantly associated bubble), keeping the alignments in the graph node coordinate system, and assessed normalized coverage over this region. We used short-read samples of six Brown Swiss cattle from which twelve haplotypes were assembled to estimate the accuracy of the pangenome alignments and coverage counting. All twelve haplotype-resolved assemblies had the deletion allele, retaining approximately 5.5 kb of the segmental duplication sequence. The mean short read coverage across the SV region predicts a sequence length closer to 8000 ± 620 bp, suggesting the coverage estimates are inflated, likely due to the abundant repetitive elements or bias from extracting reads based on alignments to the linear reference. The alignment mapping qualities were also reduced in this region, with many reads with a mapping quality of 0. Filtering out these reads removed almost all reads across this region, and so we retained these ambiguous alignments but ensured each read would correspond to only a single alignment (using the ‘--max-multimap=1’ parameter in vg giraffe).

Normalized coverage ± 1 Mb from the candidate SVs (excluding the SVs themselves) was uniform across all breeds regardless of head coloration (Figure 3A), corroborating our graph alignment and normalization approach works as intended. In contrast, the normalized coverage within the first SV region was consistently missing in white-headed breeds but inconsistently present in color-headed breeds (Figure 3B, Supplementary Figure 7), indicating poor association with the phenotype when considering additional breeds not represented in the graph. The coverage in the second, more strongly associated SV region was consistently normal in color-headed breeds and consistently elevated in white-headed breeds (Figure 3C), apart from Montbéliarde, and Normande, confirming the duplication is widely present in and likely unique to white-headed breeds.

**Figure 3.**
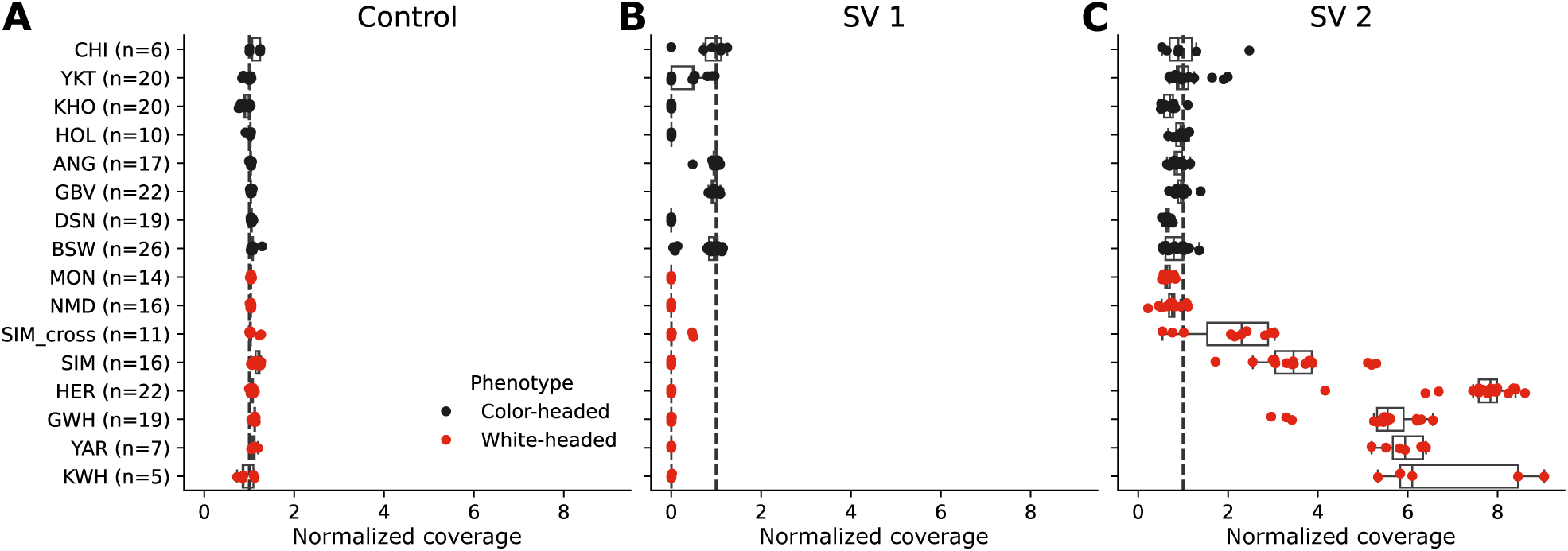
Sequencing depth in short read samples from 15 taurine breeds. (A) Normalized coverage (normalized over both sequencing depth and length of the region) per breed 1 Mb up- and downstream of the SV region. The dashed line indicates the expected normalized coverage of 1. (B) Similar to (A), but for the smaller SV 1 region. (C) Similar to (A), but for the larger SV 2 region.

Within the primary SV candidate region, Hereford animals had the highest average short-read coverage, corresponding to approximately 7-9 copies of the segmental duplication unit. Two additional Hereford samples in previously reported optical maps (Talenti et al., 2022) show an 88.5 kb insertion, corresponding to only about 6 additional copies of the 14.3 kb segmental duplication unit compared to ARS-UCD1.2, again suggesting that coverage estimates from short-read alignments may be slightly inflated. We also corroborated from older PacBio Continuous Long Read alignments that Hereford genomes most likely contain more copies of the segmental duplication than Simmental genomes (Supplementary Figure 8). The 16 Simmental samples had roughly between 3-5 copies, in agreement with the 3 copies derived from the haplotype-resolved assemblies. Samples with lower coverage could possibly indicate heterozygous carriers, as would be expected given that a small number of color-headed animals which are expected to carry two copies of a haplotype without duplication exist in the Simmental breed (Qanbari et al., 2014). Two F1s (RGVxSIM and GIRxSIM) had an intermediate increase in coverage, consistent with one haplotype (Simmental) contributing multiple copies of the repeat unit and the other haplotype (color headed) contributing only part of a copy. This was also broadly true for 7 Holstein x Simmental crosses, although some appeared to inherit the color-headed allele from Simmental parents with fewer copies.

Given the variability within the group of white-headed cattle breeds studied, including the decreased coverage in Montbéliarde and Normande, we examined the specific coverage patterns in node space (not reference coordinate space) per breed (Supplementary Figure 9). As expected, we find color-headed breeds have roughly uniform normalized coverage of 1 across the expected allele path, with little coverage outside (Figure 4A). However, Montbéliarde and Normande show elevated coverage in three parts of the white-headed allele path (Figure 4B), suggesting they may contain multiple partial copies of the segmental duplication, or a different allele structure not captured by the assemblies contained in the pangenome graph. Again as expected, the white-headed breeds with overall elevated coverage had fairly uniform coverage across different nodes (Figure 4C), primarily differing in the suspected number of copies rather than suggesting different allele structures. The Hereford samples also contained even coverage at the 1.8 kb and 800 bp sequences not observed in the ARS-UCD1.2 reference genome, suggesting that white-headed Hereford cattle contain copies of the segmental duplication matching those observed in the Simmental assemblies, but the reference genome is misassembled at this locus in terms of both copy content and number. The nodes with elevated coverage in both Montbéliarde and Normande and the other white-headed breeds correspond to regions rich with L1MAB (non-LTR retrotransposon) and MER45B (DNA transposon) repeats, possibly indicating that these elements are functionally relevant sequence for the phenotype.

**Figure 4.**
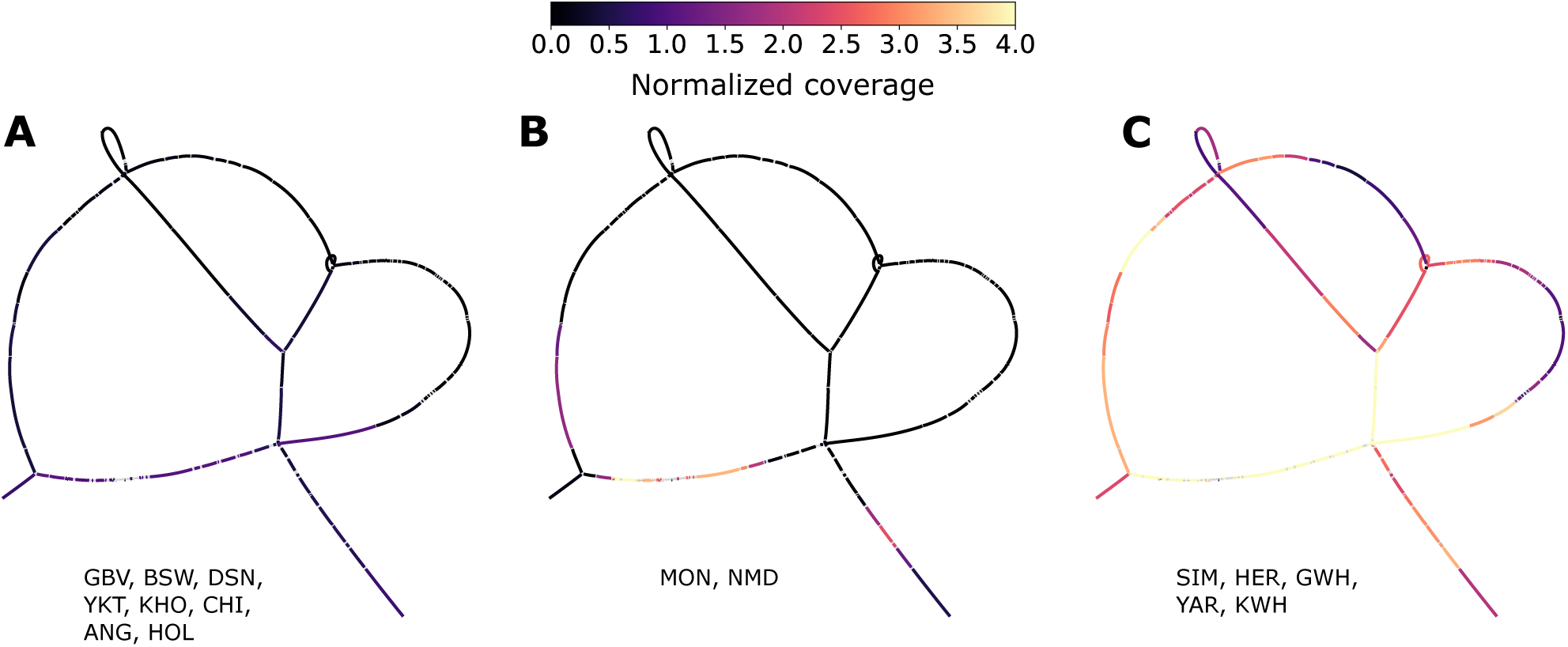
Normalized graph coverage for the three observed coverage patterns. Coverage is averaged across all samples for each node for (A) GBV, BSW, DSN, YKT, KHO, CHI, ANG, and HOL, (B) MON and NMD, and (C) SIM, HER, GWH, YAR, and KWH. Lighter node colors indicate more aligned coverage to that node.

Calculating normalized coverage over pre-identified regions is a powerful approach to estimate copy number, but requires knowledge of the phenotype status of the assemblies used to construct the pangenome to correctly partition nodes. We also investigated whether we can leverage the graph for phenotype-associations without preselecting regions to evaluate. Akin to the Jaccard similarity approach, we considered the ratio of per-node coverage between the 250 white- and color-headed short read sequencing samples. We observed a strong spike in coverage for nodes located within the candidate SV 2 region (Supplementary Figure 10), while the ratio in the discounted SV 1 region was not significant, mirroring the coverage pattern obtained for the subgraph (Figure 3). To test for any bias in the subgraph extraction/alignment process, we also combined each autosome graph into a whole-genome graph containing 51 million nodes. After realigning short read sequencing for eight samples (four of the most extreme samples from each phenotype state), we identified a clear peak in a region later confirmed as being located within the SV 2 bubble (Supplementary Figure 10).

We conducted an association study with 841,652 SNPs and Indels located on chromosome 6 that were genotyped from reference-guided short read alignments of 729 and 210 animals from the eight colored and seven white-headed breeds to investigate if a multi-breed association study can recover the head pigmentation association which we established from the graph alignment coverage pattern. Despite explicitly modeling relationships through a kinship matrix, the inclusion of individuals with distinct ancestries and the varying sample sizes across breeds likely contributed to a substantial inflation of the test statistic. Nevertheless, this analysis revealed a strong association signal upstream of the *KIT* gene (Supplementary Figure 11). The two most significantly associated SNPs (Chr6:70103450 with p=2.5e-82 and Chr6:70103502 with 1.2e-78) were over 10 orders of magnitude more significant than any other markers. Both SNPs were within the sequence that is duplicated in white-headed animals. There were only three variants overlapping coding sequence of *KIT*. All three were nonsynonymous variants and exhibited substantially higher p-values (p>9.2e-09) than the top variant further supporting that coding variants are unlikely to underly this type of head depigmentation.

## Discussion

Alleles of a segmental duplication upstream of *KIT* are associated with the characteristic white heads of Simmental and Hereford cattle, two globally occurring breeds of cattle known for dominant inherited white spotting patterns. Two F1s (RGVxSIM, GIRxSIM) originating from the crossing between white-headed Simmental cows and color-headed sires from the Gir and Rätisches Grauvieh breeds both have white heads (Figure 1A, Supplementary Figure 12) and carry one copy of the haplotype containing the segmental duplication (SIM_2, SIM_3) which supports a previously reported dominant inheritance of the white-headed phenotype (Ibsen, 1933; Qanbari et al., 2014). The white head is a breed-defining pattern in Simmental cattle where most of or all the head is depigmented against a pigmented body with white legs and underside, in addition to varying amounts and extent of white body spots as seen in other breeds such as Holstein and Montbéliarde. A typical Hereford animal is characterized by a white head, a white stripe running from the back of the head to the withers, and white legs and underparts (Supplementary Figure 13). Evidence for a series of multiple alleles at the so-called spotting locus was reported already more than 90 years ago, explaining the different degree of white pattern in Simmental and Hereford cattle (Ibsen, 1933). A potential negative side-effect of white-headed cattle, similar to gonadal hypoplasia in *KIT*-related color-sided cattle (Venhoranta et al., 2013), is an increased susceptibility to infectious bovine keratoconjunctivitis, eye cancer, or squamous cell carcinoma under UV light exposure (Anderson, 1991). However, a peculiar pigmentation around the eyes, which is a moderately to highly heritable trait associated with reduced susceptibility to eye disease, occurs in some white-headed cattle breeds (e.g., Fleckvieh, Simmental, Hereford, Groningen White Headed) and *KIT* alleles possibly contribute to this pigmentation pattern (Gonzalez-Prendes et al., 2022; Jara et al., 2022; Pausch et al., 2012).

The visible white markings are caused by a lack of melanocytes in the hair follicles and skin. We suspect that the dominance of the different white-head associated alleles reflects a gain of function resulting from dysregulated expression of the affected *KIT* gene. We thus provide another example of how iconic coat color phenotypes, which are common in domestic animal breeds, may serve as an excellent system for identifying mutations that affect fundamental processes of melanocyte development, migration, survival, and proliferation. Such mechanisms have frequently been described for *KIT* and *ASIP* mediated depigmentation patterns observed in several species (Artesi et al., 2020; Bannasch et al., 2021; Brooks et al., 2008; David et al., 2014; Henkel et al., 2019). Our analyses suggest that alleles of a segmental duplication upstream the *KIT* gene impact head pigmentation through a regulatory mechanism as the tandemly duplicated sequences do not overlap coding sequence. The repeat elements identified within the duplication may have regulatory function themselves or the breakpoints of the inserted sequence may overlap regulatory elements, thereby impacting the expression of *KIT* which leads to depigmented heads. Lifting over human genome annotations to the cattle reference localized several putative enhancer sequences nearby the segmental duplication which may be involved in regulating *KIT* expression but none of the predicted regulatory sequences was disrupted by the duplication. However, this approach neglects bovine-specific regulatory elements and is unable to correctly resolve the coordinates of diverged regulatory sequences, requiring further validation that the predicted enhancer elements exist and are relevant in the bovine genome. It could be speculated that the insertion of a largely repetitive sequence itself has no immediate effect but moves a distant enhancer of *KIT* even further upstream. Such a positional separation between regulatory and coding sequences could potentially affect the spatiotemporal expression of *KIT* during development (Berrozpe et al., 2013, 2006, 1999), thereby causing depigmentation of the head in animals carrying at least one copy of the duplication. Additional copies of the segmental duplication would further increase the distance between the enhancer sequences and *KIT*, potentially further dysregulating its expression. Such a mechanism could explain the larger area of depigmentation in Hereford cattle than in Simmental cattle, and the partial phenotypes observed in Montbéliarde and Normande (which more commonly have colored markings around the eyes/cheeks/muzzle), which likely contain multiple partial copies of the duplication. The discovery of serial alleles associated with head pigmentation expands a growing body of research on the role of allelic heterogeneity at the *KIT* locus in determining depigmentation phenotypes in large mammals such as horses (Haase et al., 2007), cattle (Artesi et al., 2020; Durkin et al., 2012; Küttel et al., 2019) and pigs (Fontanesi, 2022; Rubin et al., 2012).

We also observe that the colored-head associated “deletion” allele (with respect to the Hereford-based ARS-UCD1.2 reference) is extremely common and therefore most likely represents the ancestral bovine *KIT* allele. In addition to the 12 color-headed taurine breeds examined here, we identified the same 20.6 kb deletion in optical mapping data from six Sanga (African taurine x indicine) cattle breeds (Talenti et al., 2022), as well as in pangenomes containing three further taurine breeds (Jang et al., 2023) and two indicine breeds and three non-cattle bovids (Leonard et al., 2023a) (Supplementary Table 3). Thus, it is very likely that the color-headed phenotype is the ancestral state, while the white-headed phenotype has evolved by mutation(s). Although there are no assemblies to confirm the allelic structure, the Russian Yaroslavl breed also appears to have a similar SV to Hereford and Simmental, despite geographically limited opportunities of introgression (Zinovieva et al., 2020). However, there is limited metadata and no pedigree available to confirm the ancestry of the public Yaroslavl samples. The smaller 6-7 kb SV is also widespread but again is less consistent with the color-headed phenotype, suggesting that it does not play a role in head pigmentation or may affect a different phenotype.

The number of copies of the segmental duplication also varies within breeds. We observed considerable variation in Simmental and Fleckvieh cattle, as expected given the sporadic occurrence of color-headed individuals in these breeds (Qanbari et al., 2014). One can speculate that the degree of depigmentation is dosage-dependent or that the segmental duplication contributes to depigmentation beyond the head. However, such investigations require extensive phenotype observations and precise copy number resolution, neither of which are available for the public short-read samples considered in our study. The presence of other *KIT* alleles associated with head and coat color phenotypes (Hayes et al., 2010; Pausch et al., 2012) may further complicate such an analysis.

The use of pangenomes for association testing has largely relied on presence/absence variation between groups (Brynildsrud et al., 2016; Leonard et al., 2022), decomposed graph genome variation (Chin et al., 2023), or genotypes obtained by mapping of short reads to pangenomes (Cochetel et al., 2023; Sirén et al., 2021). We have developed and applied an approach that compares path similarities and differences between assemblies of samples with divergent phenotypes.

While this pangenome “phenotype signature” analysis successfully identified and resolved a promising SV associated with a white-headed phenotype, the Jaccard similarity metric is less sensitive to small variation, especially given that splitting a graph into bins much smaller than 1 kb may completely disrupt the graph topology. However, compared to a GWAS or other genotype-based tests that can be conducted from decomposed or short read-called graph genome variation with standard tools (e.g., (Yang et al., 2011)), the Jaccard similarity metric is more flexible for testing multiallelic variation, as the relevant paths can have high similarity even if the genotypes are distinct. Furthermore, non-reference bias is only mitigated for the breeds represented in our pangenome; any additional, unrepresented allelic structure (e.g., possibly in Montbéliarde and Normande) is still likely lost in downstream alignment and analysis. Potential misassemblies, like that likely found in the reference genome, as well as suboptimal or redundant graph topology further complicate association testing on graphs. We also demonstrate that phenotype associations can be identified from a whole-genome graph without first detecting a candidate region based on assembly phenotype status. This would allow a single diverse-but-generic pangenome to be used for testing any traits for which there were sufficient sequenced samples.

As assemblies become increasingly available across almost all species, and in particular for bovines (Smith et al., 2023), the approaches outlined here will become increasingly powerful for resolving complex variation associated with binary phenotypes. These approaches could also be generalized to quantitative traits with a more refined calculation of coverage across nodes. This will be particularly important in cases such as those presented here where even long read alignments to a linear reference fail to consistently resolve the variation. However, such a pangenome-wide association mapping approach requires further refinement to correct for confounding factors such as population stratification and hidden relatedness which can cause spurious associations (Price et al., 2006; Sul et al., 2018).

## Methods

### Ethics statement

The sampling of blood was approved by the veterinary office of the Canton of Zurich (animal experimentation permit ZH 200/19).

### Animals

Two purebred Simmental (*Bos taurus taurus*) cows were inseminated with semen from Gir (*Bos taurus indicus*) and Rätisches Grauvieh (*Bos taurus taurus*) sires. A purebred Evolèner cow was inseminated with semen from a dwarf zebu (DWZ). Two female calves (GIR x SIM and DWZ x EVO) and a male calf (RGV x SIM) were delivered at term. DNA was prepared from blood samples of the F1s and dams, and from cryopreserved semen samples of the sires. DNA of a purebred Brown Swiss cow was prepared from a blood sample provided by Braunvieh Schweiz. DNA of three Braunvieh (cross between Brown Swiss and Original Braunvieh) and one Original Braunvieh bulls was prepared from testis tissue sampled at a commercial slaughterhouse from a previous project (Mapel et al., 2024).

### Sequencing

PacBio HiFi reads were collected from four F1s used for trio-binning with three SMRT Cell 8M sequenced for each sample on a Sequel IIe, and from another four animals used for dual-assembly with a total of four SMRTs Cell 25M sequenced on a Revio and five SMRT Cells 8M sequenced on a Sequel IIe. Illumina paired-end (2x150 bp) reads were collected from the four F1s and their parents.

### Genome assemblies

The Hereford (HER), Simmental (SIM_1), Highland (HIG), Original Braunvieh (OBV_1 and OBV_2), and Brown Swiss (BSW_1, BSW_2, BSW_5, BSW_6, and BSW_7) were downloaded from public sources (Supplementary Table 1). We assembled the new genomes with hifiasm (v0.19.5) (Cheng et al., 2021) with default parameters. Where available, we used parental short reads to create haplotype-resolved assemblies with yak (v0.1). Contigs were then scaffolded to the ARS-UCD1.2+Y reference with RagTag (v2.1.0) (Alonge et al., 2022). We assessed assembly quality with merqury (6b5405) (Rhie et al., 2020) and completeness with compleasm (v0.2.2) (Huang and Li, 2023) using the Cetartiodactyla lineage.

### Identification of a missing copy of the segmental duplication in the SIM_2 assembly

The SIM_2 haplotype was initially assembled with only two copies of the segmental duplication. We used the parental short reads to triobin the HiFi reads using meryl (v1.3) and Canu (v2.2) (Koren et al., 2018) before aligning to the reference genome with minimap2 (v2.24) (Li, 2018). We then extracted all HiFi reads corresponding to the Simmental haplotype from the region 6:69099582-71123136 with SAMtools fastq (v1.16.1) (Danecek et al., 2021). We reassembled these reads with hifiasm with the additional parameters -D 10 -N 500 to improve repeat sensitivity. We then manually split the original Simmental assembly between the two repeat copies and used RagTag patch to “gap fill” with the sequence of the newly assembled third copy.

### Pangenome construction

The pangenome graph was built with pggb (736c50d) (Garrison et al., 2023), using wfmash (67ab187) (Marco-Sola et al., 2021), seqwish (f44b402) (Garrison and Guarracino, 2023), smoothxg (31d99f2) (https://github.com/pangenome/smoothxg), gfaffix (0.1.5) (https://github.com/marschall-lab/GFAffix), and odgi (de70fcda) (Guarracino et al., 2022). We estimated the pangenome percent-identity with mash (v2.3) (Ondov et al., 2019, 2016) triangle as 97-99% across different autosomes, as well as a segment-length of 75k and a min-match-length of 31. We used the hierarchy linkage clustering with the UPGMA algorithm from scipy (v1.11.4) (Virtanen et al., 2020) to approximate the phylogenetic tree across chromosome 1 after removing centromeres, which are unevenly present on assemblies. We then extracted the relevant subgraph using odgi extract for the region 6:69099582-71123136 followed by cycle-breaking with odgi sort -p “wc”. The subgraph region was visualized with BandageNG (v2022.09) (https://github.com/asl/BandageNG) (Wick et al., 2015).

We then built a whole-genome pangenome using odgi squeeze to combine all 29 per-autosomes graphs into one large graph, renaming nodes and paths as necessary to remain unique. We then trimmed centromeric regions (as identified per-assembly) in all non-reference samples using odgi extract --inverse --bed-file <centromere regions> and collapsed runs of N nucleotides with odgi crush. We then further simplified the graph using vg (v1.51.0) (Sirén et al., 2021) clip -d 3 -m 10000 while retaining all reference nodes in order to produce a graph which could be indexed.

### Calculation of the Jaccard similarity ratio

We used odgi extract to bin every 1 kb of reference sequence into subgraphs, and then used odgi similarity to calculate the Jaccard similarity of all paths across these subgraphs. We calculated the Jaccard ratio as the ratio of intra-group similarities (white-white or colored-colored) to inter-group similarities (white-colored), such that a high value demonstrates both groups are internally consistent but distinct across groups.

### Identification of repetitive elements in the graph bubble

We used RepeatMasker (https://www.repeatmasker.org/, v4.1.5) with the repbase repeat library to identify repetitive elements in the SIM_3 assembly haplotype. We used odgi procbed to adjust the repeat coordinates into the subregion spanned by the graph bubble, before using odgi inject to annotate the graph with repeat paths. Finally, we used odgi untangle with “-g” to produce a sequence arrow map, which was then visualized in R with the packages gggenes (v.0.5.1) and ggplot2 (v3.4.4).

### Short read data collection and mapping to the graph

We downloaded publicly available short read sequencing data from study accessions PRJNA814817, PRJEB9343, PRJNA176557, PRJEB45822, PRJEB56301, PRJNA494431, PRJNA762180, PRJNA642008 and PRJEB18113 (Supplementary Table 1) using fastq-dl (v2.0.4) with “--group-by-sample”. All fastq sequences were trimmed with fastp (v0.23.4) (Chen et al., 2018) using default parameters. Reads were then aligned to ARS-UCD1.2+Y using strobealign (6bbc5b7) (Sahlin, 2022), followed by read group sorting, duplicate marking, and coordinate sorting with SAMtools. We then extracted reads from the region of interest with SAMtools fastq, and then aligned these reads to the pangenome using vg giraffe with “--named-coordinates --max-multimaps 1 -o gaf”. For the whole-genome alignments, we aligned the entire trimmed fastq to the whole-genome graph without any selection of reads. We assessed per-node coverage with gafpack (v0.1.0).

### PCA breed validation

Variants were called from the aligned BAM files using DeepVariant (v1.6) (Poplin et al., 2018), using the WGS model on chromosomes 1-3. We conducted a PCA using plink2 (v2.00a4LM) (Chang et al., 2015), first thinning by linkage above r>0.8 every 100 kb, excluding variants below minor allele frequency of 10%, and only retaining biallelic SNPs. Samples were then colored by assigned breed, confirming the accuracy of the metadata associated with each SRA sample.

### Detecting signatures of selection

We used sequence variant genotypes from run 9 of the 1000 Bull Genomes project (Hayes and Daetwyler, 2019) for 49 SIM/FV, 50 BSW, and 33 HER cattle to identify signatures of selection. Accession numbers of the raw sequencing data are in Supplementary Table 2. We retained genotypes for 2,347,081 SNPs located on chromosome 6 for which Dorji et al. (Dorji et al., 2024) assigned the ancestral allele with a probability greater than 0.85. Using a pre-computed empirical allele frequency spectrum of the chromosome 6 SNPs, we calculated composite likelihood ratio (CLR) statistics in non-overlapping 50 kb windows using SweepFinder2 (v1.0) (DeGiorgio et al., 2016).

### Small variant association testing

We considered sequence variant genotypes from run 9 of the 1000 Bull Genomes project (Hayes and Daetwyler, 2019) for 729 and 210 animals from eight colored (CHI, KOL, YAK, HOL, ANG, GBV, BSW, DSN) and seven white-headed (SIM, GWH, HER, MON, NOR, YAR, KWH) breeds. Accession numbers of the raw sequencing data are in Supplementary Table 2. We considered all autosomal variants with minor allele frequency greater than 1% to build a genomic relationship matrix with GCTA (v 1.92.1) (Yang et al., 2011). The “--mlma”-option of GCTA was subsequently used to conduct a mixed model-based association analysis with all variants located on chromosome 6.

### Validation of the SV in other breeds and species

We called SVs from each of the aligned HiFi samples over the region of interest using sniffles2 (v2.2) (Smolka et al., 2024) before merging the resulting snf files into a joint-called VCF.

We used the merged VCF file of structural variants called from optical mapping data from (Talenti et al., 2022) to investigate the presence of the candidate SV in taurine and indicine cattle. We also examined the 14-breed pangenome from (Jang et al., 2023), first realigning the constituent assemblies to the pangenome using minigraph --call (v0.20) (Li et al., 2020), followed by examining the paths taken through bubble variation as described in (Leonard et al., 2023a). We also aligned PacBio Continuous Long Read data from the Simmental, Holstein, Hereford, Original Braunvieh and Evolèner breeds available at the study accession PRJEB72196 to the reference genome using minimap2 with the “map-pb” preset. We then extracted all aligned reads over the region BTA6:69099582-71123136 using SAMtools fastq, and then used minimap2 to align the 14.3 kb segmental duplication to these reads, counting the number of hits > 10 kb to estimate the copy number per sample.

### Prediction of regulatory elements overlapping the segmental duplication

We downloaded coordinates from publicly available human regulatory elements curated by the FANTOM5 (Andersson et al., 2014) and ENCODE (Moore et al., 2020) consortia from https://fantom.gsc.riken.jp/5/datafiles/latest/extra/Enhancers/human_permissive_enhancers_phase_1_and_2.bed.gz and https://downloads.wenglab.org/V3/GRCh38-cCREs.bed, respectively. Genome coordinates of human regulatory elements were converted from the GRCh37/hg19 and GRCh38/hg38 assemblies to the ARS-UCD1.2 assembly using the LiftOver tool (https://genome.ucsc.edu/cgi-bin/hgLiftOver) from the UCSC Genome Browser.

## Data Access

HiFi reads used for genome assembly are available in the European Nucleotide Archive (ENA) (https://www.ebi.ac.uk/ena/browser/home) at the study accession PRJEB42335 under sample accessions SAMEA115129361, SAMEA113612082, SAMEA113612088, SAMEA113612091, SAMEA115129362, SAMEA115129363, SAMEA115129364, and SAMEA115129365. Parental and F1 short reads are available in the ENA at the study accession PRJEB28191 under sample accessions SAMEA115121766 and SAMEA115121767 for the RGVxSIM (SAMEA115129365), SAMEA115121768 and SAMEA115121769 for the DWZxEVO (SAMEA115129364), SAMEA115121770 and SAMEA115121771 for the GIRxSIM (SAMEA115129363), and SAMEA115121772 and SAMEA115121773 for the BSWxBSW (SAMEA115129361). PacBio CLR data are available at the study accession PRJEB72196 under sample accessions SAMEA115160498, SAMEA115160497, SAMEA115160496, SAMEA115160495, SAMEA115160494, and SAMEA115160493. All scripts and workflows are available online (https://github.com/AnimalGenomicsETH/pangenome_KIT).

## Acknowledgements

We are thankful for the technical support provided by Dr. Anna Bratus-Neuenschwander from the ETH Zürich technology platform Functional Genomics Center Zurich (https://fgcz.ch) and the Next Generation Sequencing Platform at the University of Bern for sequencing and DNA fragment analysis. We also thank Eirini Lampraki from Pacific Biosciences for DNA sequencing on a Revio system. We thank Flavio Ferrari (AgroVet-Strickhof) for animal handling.

## Author contributions

SM analyzed pangenome variation, interpreted results, and wrote the manuscript; ASL assembled and aligned long reads, developed the phenotype signature approach, analyzed pangenome variation, analyzed short-read data, interpreted results, and wrote the manuscript; XMM extracted high molecular weight DNA and revised the manuscript; SMBU was responsible for animal experimentation and the sampling of blood; CD contributed data, interpreted results and contributed to the writing of the manuscript; HP conceived the study, interpreted results, and wrote the manuscript; all authors approved the final version of the manuscript.

