## Supporting Tables and Figures for "Taurine pangenome uncovers a segmental duplication upstream of *KIT* associated with depigmentation in white-headed cattle": Supplementary_Figures_and_Tables.docx

**Supplementary Material**

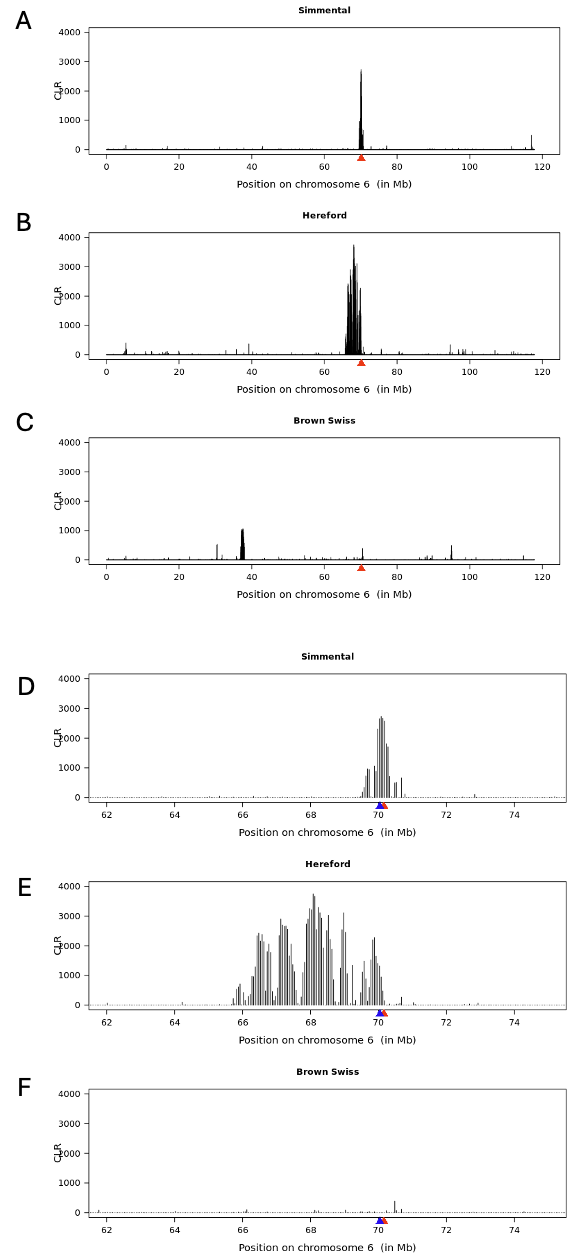

Supplementary Figure 1: Signatures of selection detected in different breeds of cattle. (a-c) Composite likelihood ratio (CLR) of 50kb windows on bovine chromosome 6 for Simmental (a), Hereford (b), and Brown Swiss (c). The red triangle indicates the position of the KIT gene. (d-f) same as in (a-c) but zoomed into a 15 Mb window centered on KIT. The blue triangle indicates the position of the SV identified in the pangenome analysis.

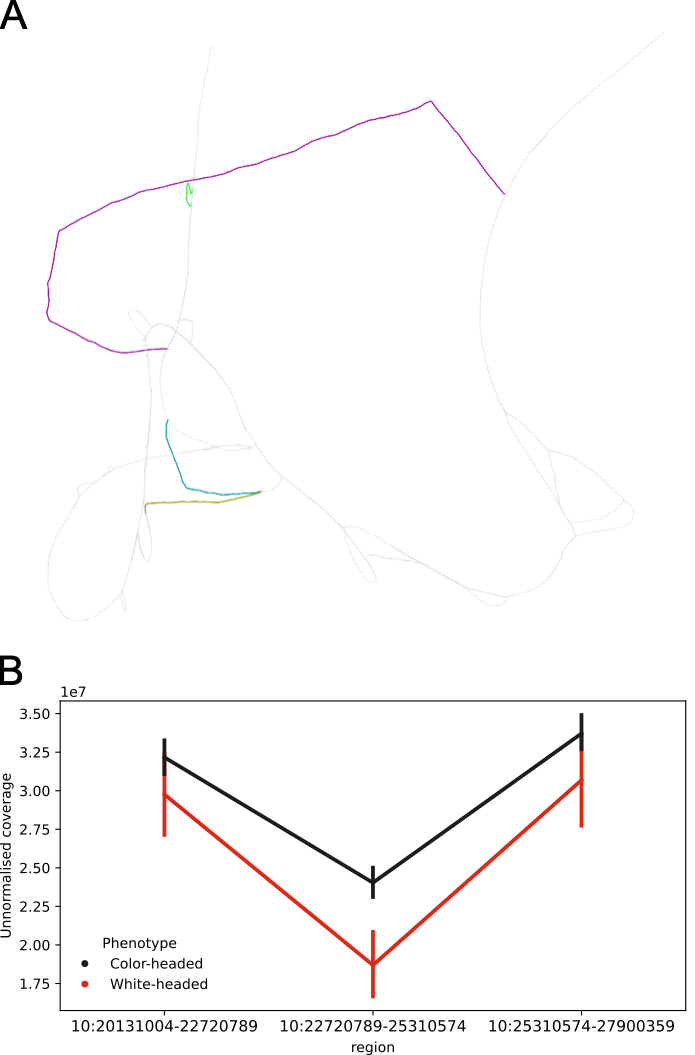

**Supplementary Figure 2**. (A) Poorly constructed region leads to inflation of non-reference sequence. Large regions in ARS_UCD1.2 (pink, 2.6 Mb), a Simmental (yellow, 602 kb), and two Brown Swiss (blue, 410 kb and green, 402 kb) shared homology with other nodes, but were not successfully “squished” into the graph during the seqwish or smoothxg steps of graph building due to suboptimal alignments. (B) Unnormalised coverage (raw number of bases aligned) of 251 short read sequencing samples aligned to the ARS-UCD1.2 reference genome show a clear drop in coverage in the region identified in (A) with relatively stable coverage in a similarly sized window before and after the region.

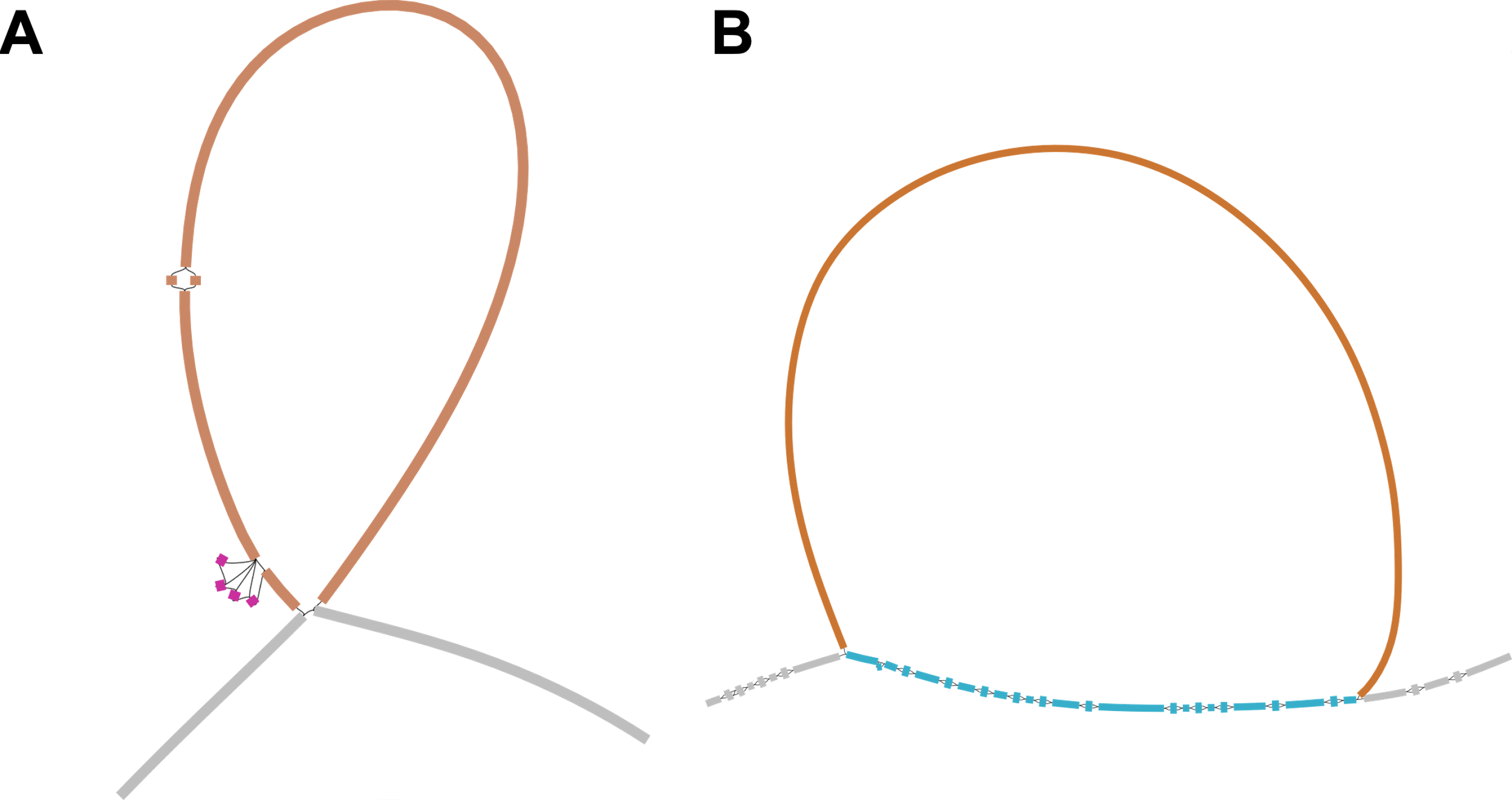

**Supplementary Figure 3**. Detailed view on regions with nearly significant Jaccard similarity on (A) BTA6:54.7 Mb and (B) BTA21:47.3 Mb. (A) The three Simmental and the Hereford assemblies took the orange 1.2 kb insertion (annotated as an L1 repeat), but so did one Brown Swiss and the Highland assembly. The small pink nodes were mostly unique to Simmental. (B) The three Simmental and the Hereford assemblies looped through the blue nodes a second time after taking the orange 2.4 kb insertion (annotated as BovB RTEs and ERV1 repeats), but so did the Evolener and the Highland assemblies. All other assemblies only traversed the blue nodes a single time.

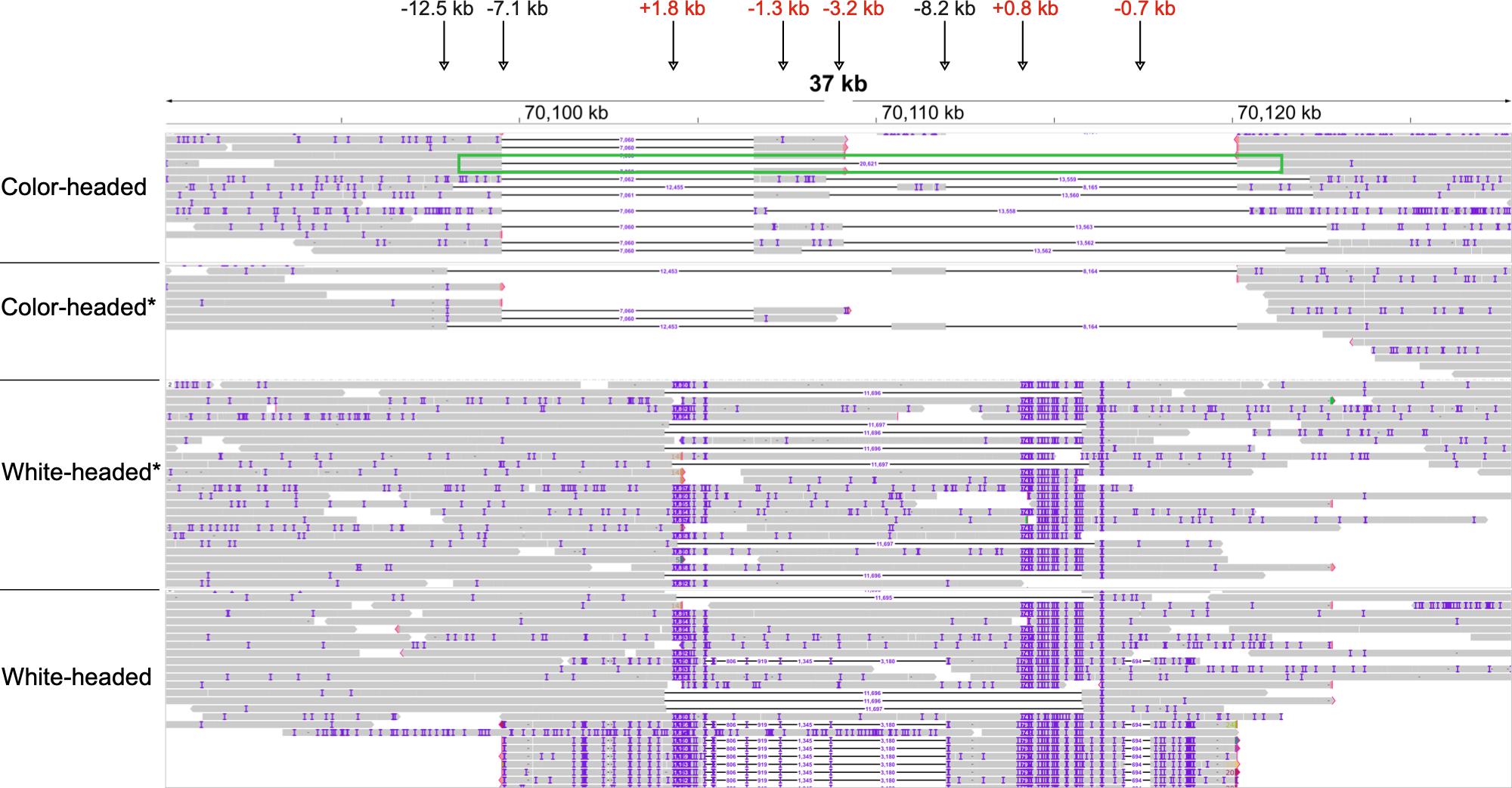

**Supplementary Figure 4:** Long read alignments produce inconsistent SV genotyping. Due to the internal repetitive structure of the duplication, even long read alignments struggle to resolve the region. Only a single read from a color-headed sample correctly aligned with the expected ~20.6 kb deletion (green box). Structural variant calling with sniffles2 reported 12.5, 7.1, and 8.2 kb deletions for color-headed samples (black SVs) without a clear pattern of allele assignment. Conversely, no reads for white-headed samples corrected aligned the expected ~16 kb insertion. Instead, sniffles2 only reported 1.8 and 0.8 kb insertions (correctly matching the known SIM-specific bubbles in the graph) but also 1.3, 3.2, and 0.7 kb deletions for white-headed samples (red SVs).

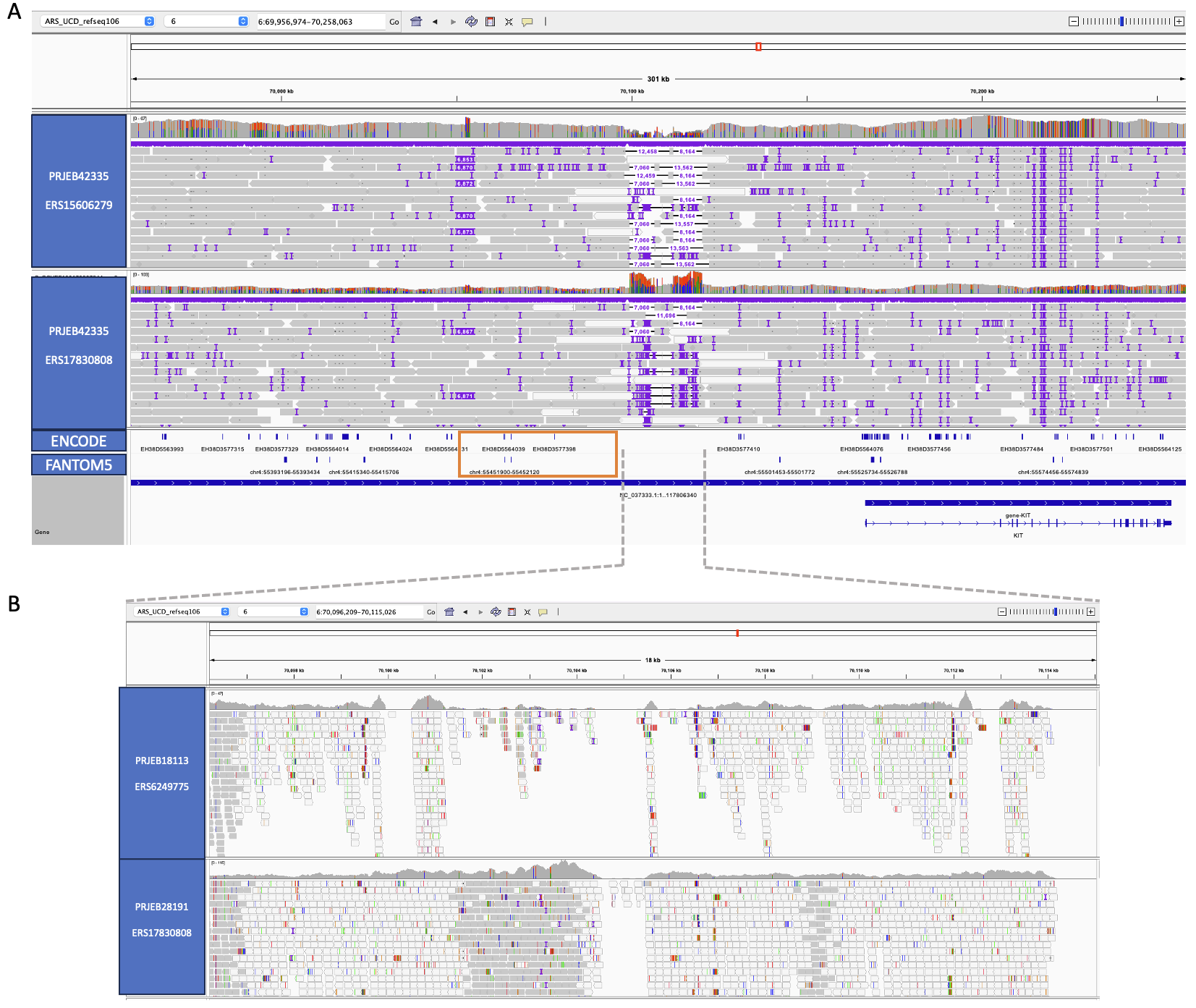

**Supplementary Figure 5**. IGV screenshot of the candidate SV region. A) HiFi reads aligned against ARS-UCD1.2 for two samples (ERS15606279: BSW x BSW; ERS17830808: GIR x SIM) indicate several large structural variants upstream KIT. Putative cis-regulatory elements were lifted over from FANTOM5 and ENCODE. The orange box contains four candidate cis-regulatory elements (EH38D3577398 EH38D3577384, EH38D3577379, EH38D5564039) predicted by human ENCODE which were lifted from the human reference genome including two that also overlapped lifted coordinates of permissive enhancers predicted by FANTOM5. B) Illumina short paired end read alignments for a subset of the region (IGV can not visualize the full region for short reads) for the same animals (ERS6249775: BSW x BSW; ERS17830808: GIR x SIM). The associated SV bubble is impossible to resolve with short reads.

**Supplementary Figure 6**. Top two principal components of SNPs called from 250 short read sequencing samples. Colors indicate breed metadata associated with each sample.

**Supplementary Figure 7**. Normalized coverage (normalized over both sequencing depth and length of the region) per breed over the reference path within the SV 1 region which all breeds take. The dashed line indicates the expected normalized coverage of 1.

**Supplementary Figure 8**. CLR alignments from a 200 kb window around the KIT bubble, measuring significant (longer than 2.5 kb) alignments to the segmental duplication sequence. Color-headed animals (Eringer, Original Braunvieh, and Holstein) have no support for more than 2 hits (measuring the expected two hits on either end of the graph bubble), while one Hereford and two Simmental samples have clear support for additional copies. There are not sufficient ultralong reads (> 56 kb) to genotype more than 4 copies in any single read, as predicted for Hereford samples.

**Supplementary Figure 9**. Normalized coverage per node per breed for (A) color-headed and (B&C) two patterns of white-headed phenotypes. Although MON and NMD have an overall average coverage similar to color-headed breeds in Figure 3, they have elevated coverage in some regions similar to other white-headed breeds followed by regions of zero coverage.

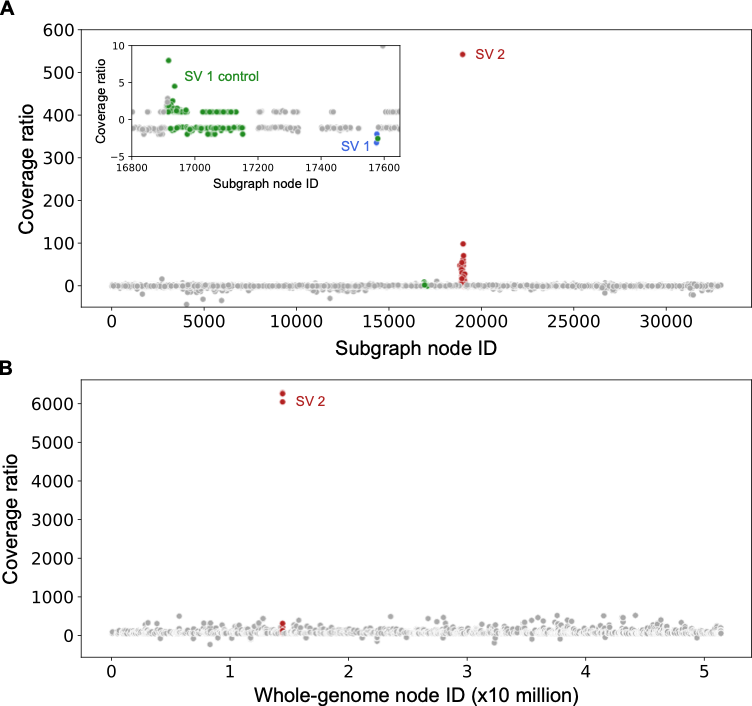

**Supplementary Figure 10**. Graph-based phenotype association test. (A) Taking the ratio of the median white-headed to color-headed coverage (or the negative reciprocal if the ratio is less than one) for the 250 samples reveals a clear peak associated with nodes identified as part of SV 2 (the candidate region). The nodes identified as part of SV 1 or its reference path (SV 1 control) were not significantly associated with the phenotype, as seen in Figure 3. (B) Whole-genome alignment for a subset of eight samples (four of the most extreme samples for each phenotype state based on the subgraph analysis) also reveals a clear spike in nodes identified as part of SV 2. The SV 1 regions are not colored for simplicity.

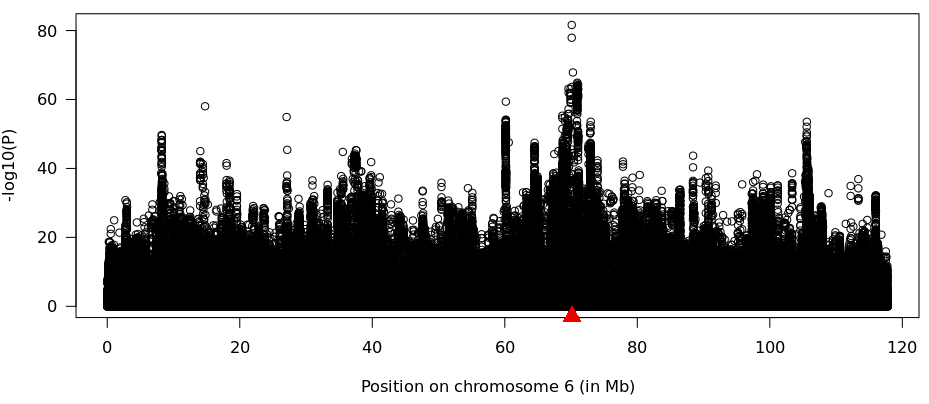

**Supplementary Figure 11**. Association study with sequence variants of 729 and 210 animals from eight colored and seven white-headed breeds. The red triangle indicates the position of KIT.

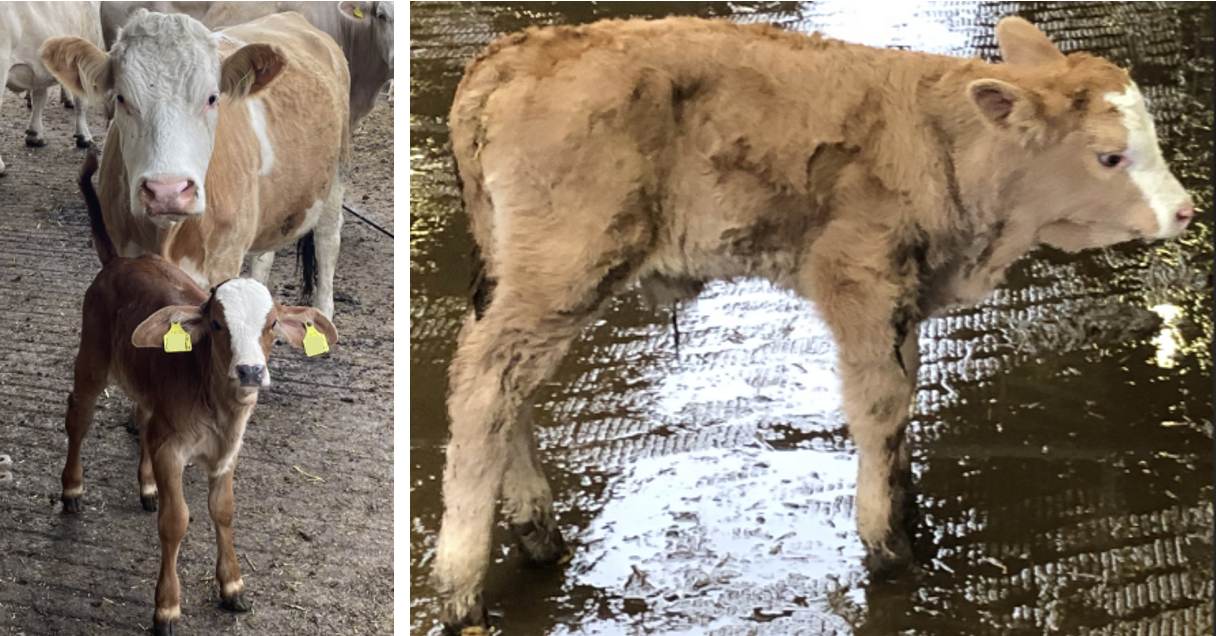

**Supplementary Figure 12.**. Head depigmentation in an RGVxSIM F1 calf.

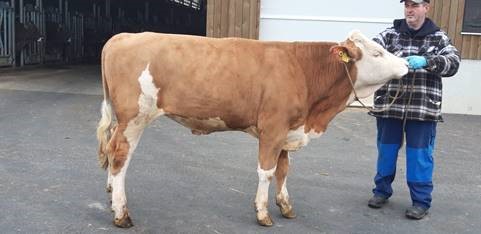

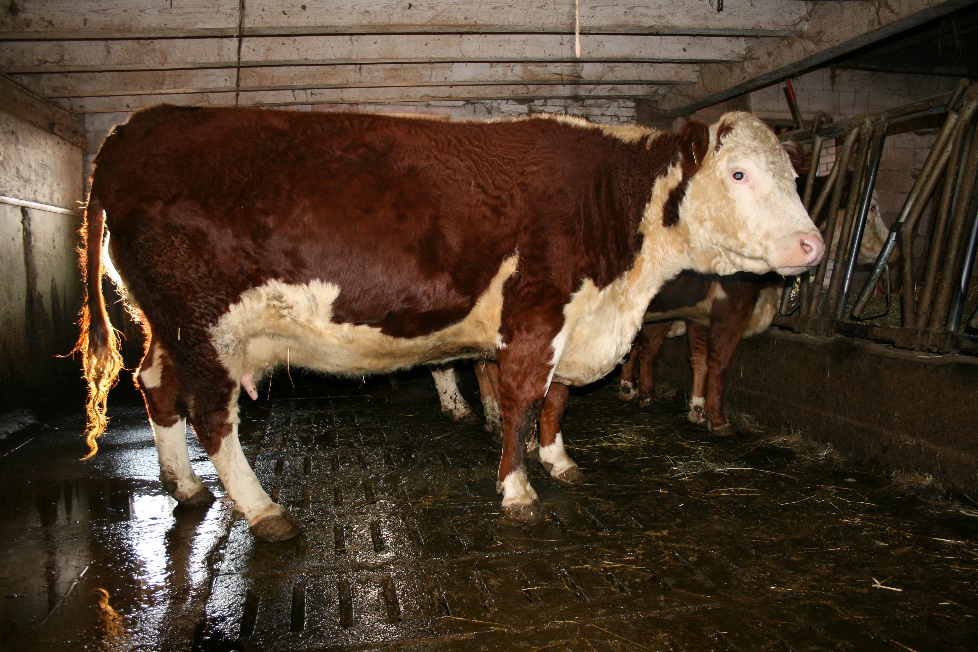

**Supplementary Figure 13**. Pigmentation patterns in Simmental and Hereford animals. A) A typical male Simmental bull characterized by a white face, and white legs and white underparts. B) A typical Hereford cow characterized by a white face, a white stripe running from the back of the head to the withers, and distally white legs and large white underparts. Note the extensive area of continuous depigmentation in the neck and at the withers.

**Supplementary Table 1:** Previously published assemblies that were integrated into the pangenome graph. CLR: PacBio Continuous Long Reads, HiFi: PacBio HiFi reads, ONT: Oxford Nanopore Technologies reads.

| Sample / Acronym | Primary sequencing technology | Reference | Coat Colour Phenotype  (head) |
| --- | --- | --- | --- |
| Hereford / HER | CLR | (Rosen et al., 2020) | white-headed |
| Simmental / SIM_1 | ONT | (Heaton et al., 2021) | white-headed |
| Highland / HIG | ONT | (Rice et al., 2020) | colored head |
| Brown Swiss / BSW_1 | HiFi | (Leonard et al., 2023b) | colored head |
| Brown Swiss / BSW_2 | HiFi | (Leonard et al., 2023b) | colored head |
| Brown Swiss / BSW_5 | HiFi | (Leonard et al., 2023b) | colored head |
| Brown Swiss / BSW_6 | HiFi | (Leonard et al., 2023b) | colored head |
| Brown Swiss / BSW_7 | HiFi | (Leonard et al., 2022) | colored head |
| Original Braunvieh / OBV_1 | HiFi | (Leonard et al., 2022) | colored head |
| Original Braunvieh / OBV_2 | HiFi | (Leonard et al., 2022) | colored head |

**Supplementary Table 2**. Excel spreadsheet with three sheets listing accession numbers for public short read samples considered in our study.

**Pangenome mapping:** 251 taurine cattle short read samples used in pangenomic alignment. Arbitrary sample IDs were created when not publicly available. Coverage was determined across the autosomes, normalized by the reference length of 2,489,385,779 bp. Duplication rate was assessed by samtools during the markdup stage as the ratio of “DUPLICATE PAIR” to “PAIRED”.

**Signatures of selection: 139** taurine cattle short read samples used to identify signatures of selection

**GWAS: 739** taurine cattle short read samples from white-headed and colored breeds used in association mapping

<attached xlsx>

**Supplementary Table 3**. Additional evidence for the 6-7 kb insertion (SV 1) and 20.6 kb deletion (SV 2) in color-headed bovids, where - indicates the SV was not observed. Taurine/Indicine/Sanga are Bos taurus subspecies.

| **Species/breed** | **Species** | **SV 1** | **SV 2** | **Source** | **Reference** |
| --- | --- | --- | --- | --- | --- |
| Piedmontese | Taurine | 6,022 | -20,622 | Pangenome | (Leonard et al., 2023a) |
| Holstein | Taurine | - | -20,649 | Pangenome | (Jang et al., 2023) |
| Jersey | Taurine | - | -20,649 | Pangenome | (Jang et al., 2023) |
| Hanwoo | Taurine | 6,877 | -20,649 | Pangenome | (Jang et al., 2023) |
| Angoni | Sanga | - | -20,613 | Optical map | (Talenti et al., 2022) |
| Ankole | Sanga | - | -20,613 | Pangenome & Optical map | (Jang et al., 2023; Talenti et al., 2022) |
| Barotse | Sanga | 6,812/- | -20,613 | Optical map | (Talenti et al., 2022) |
| Boran | Sanga | 6,812 | -20,613 | Optical map | (Talenti et al., 2022) |
| Ndama | Sanga | 6,812 | -20,613 | Pangenome & Optical map | (Jang et al., 2023; Talenti et al., 2022) |
| White Fulani | Sanga | 6,812 | -20,613 | Optical map | (Talenti et al., 2022) |
| Brahman | Indicine | 6,876 | -20,622 | Pangenome | (Leonard et al., 2023a) |
| Nelore | Indicine | 6,812/- | -20,622 | Pangenome & Optical map | (Leonard et al., 2023a; Talenti et al., 2022) |
| Gaur | *Bos gaurus* | 6,876 | -20,622 | Pangenome | (Leonard et al., 2023a) |
| Yak | *Bos grunniens* | 6,876 | -20,622 | Pangenome | (Leonard et al., 2023a) |
| Bison | *Bison bison* | 6,876 | -20,622 | Pangenome | (Leonard et al., 2023a) |
